# Similarities between Borderline Personality Disorder and Post traumatic Stress Disorder: evidence from Resting-State Meta-Analysis

**DOI:** 10.1101/633248

**Authors:** Ali Amad, Joaquim Radua, Guillaume Vaiva, SCR Williams, Thomas Fovet

## Abstract

Borderline personality disorder (BPD) and post-traumatic stress disorder (PTSD) are common psychiatric disorders. The nature of the relationship between BPD and PTSD remains controversial, but it has been suggested that these disorders should brought closer because of their many similarities. We thus performed a quantitative meta-analysis of resting-state functional imaging to assess similarities in the brain activation across BPD and PTSD diagnostic groups.

Overlap analyses revealed decreased activation in the left and right precuneus of both BPD and PTSD groups when compared to control subjects. BPD showed significant increased, but PTSD showed decreased activation, relative to control subjects, in the anterior cingulate/paracingulate gyri and in the left superior frontal gyrus. Complementary overlap analyses on a subgroup of studies with similar sex and age distribution partially confirmed the main results as the same pattern of functional activation in the anterior cingulate and in the left superior frontal gyrus were found.

Our findings are in agreement with the hypothesis that BPD and PTSD share common neuropathological pathways.

## INTRODUCTION

Borderline personality disorder (BPD) and post-traumatic stress disorder (PTSD) are common psychiatric disorders with an estimated prevalence in the general population of 0.5–5.9% and 3% respectively (Bisson et al., 2015; Grant et al., 2008; Lenzenweger et al., 2007). These disorders are associated with high mortality and morbidity, due to suicide, frequent hospitalization, substance use, psychiatric comorbidity, considerable economic burden and poor quality of interpersonal relationships (Bisson et al., 2015; Skodol et al., 2002). The nature of the relationship between BPD and PTSD remains controversial but several arguments suggest that these disorders should brought closer because of their many similarities (Gunderson and Sabo, 1993; Lewis and Grenyer, 2009).

First, BPD and PTSD share numerous clinical features, particularly major disturbances in emotional regulation, impulse control, reality testing, interpersonal relationships and sense of identity (Gunderson and Sabo, 1993; Lewis and Grenyer, 2009). Second, the role of trauma is essential in both disorders. Indeed, the prevalence of childhood trauma is so frequently associated with BPD (either neglect (92%), sexual abuse (40%-70%) or physical abuse (25%-73%) (Zanarini et al., 2002)) that a causal relationship has been suggested (Ball and Links, 2009). BPD is often shaped in part by trauma (Gunderson and Sabo, 1993), and individuals with BPD are therefore vulnerable to develop PTSD (Golier et al., 2003). From a neurobiological perspective, BPD and PTSD are both associated with a hypothalamus–pituitary–adrenal (HPA) axis dysregulation. Notably, BPD patients displayed elevated continuous cortisol output and blunted cortisol following psychosocial challenges whereas cortisol levels are generally found decreased in PTSD patients (de Kloet et al., 2006; Drews et al., 2019; Schumacher et al., 2019; Wingenfeld et al., 2010).

Furthermore, genetic associations between *FKBP5* variants–an important functional regulator of the glucocorticoid receptor complex associated with PTSD in a large number of studies (Zannas and Binder, 2014)– and BPD diagnosis have been shown recently (Amad et al., 2019; Martín-Blanco et al., 2015). These results seem to indicate that BPD shares a genetic vulnerability with PTSD. Finally, from a neuro-functional perspective, BPD and PTSD are supposed to share the same abnormalities in fronto-limbic networks with a limbic hyperactivity and a diminished recruitment of frontal brain regions (Krause-Utz et al., 2014; Patel et al., 2012) even if this common assumption appears to not be supported by functional imaging meta-analysis in these two disorders (Amad and Radua, 2017; Wang et al., 2016).

To investigate the relationship between BPD and PTSD from a neuro-functional perspective, we performed a functional quantitative meta-analysis to compare results across diagnostic groups and to assess functional similarities that may reflect common neuropathological pathways in these disorders.

## MATERIAL AND METHODS

All brain imaging studies of the resting-state studies which have compared brain activation in patients with BPD or PTSD versus healthy controls were included in two separate meta-analyses (one for BPD and one for PTSD). To identify eligible studies, a systematic search, in respect to the PRISMA statement (Moher et al., 2009), was conducted on the Medline and ISI Web of Knowledge up to April 2018 using the following search term combinations: (1) ‘‘neuroimaging’’, ‘‘fMRI’’, ‘‘PET’’, “ALFF”, “ReHo”, “rCBF”, (2) ‘‘resting-state’’, ‘‘default network’’ and (3) the terms ‘‘borderline personality disorder’’ or “post-traumatic stress disorder”. Reviews and meta-analyses were cross-referenced to identify studies that were missed in the literature searches. Authors were contacted for unpublished data including t-maps from the original studies. Studies were only selected if they had performed a whole brain analysis and reported activation coordinates in standard space (MNI or Talairach). Also, we ensured that the same threshold throughout the whole brain was used within each included study, in order to avoid bias toward liberally thresholded brain regions. The literature search strategy is summarised in the flow chart presented in **Supplementary Figures 1 and 2**.

The functional neuroimaging meta-analyses were performed using the anisotropic effect size version of Seed-based D Mapping (formerly Signed Differential Mapping, AES-SDM, www.sdmproject.com) (Radua et al., 2014, 2012). This method allows the inclusion of both peak information (coordinates and t-values) and statistical parametric maps to create whole-brain effect size and variance maps, which are then used to perform voxel-wise random effects meta-analyses.

Peak coordinates and effect-sizes (e.g. t-values or z-scores) of statistically significant differences between patients and controls at the whole-brain level were extracted from each dataset. A standard MNI map of the resting-state differences was then separately recreated for each dataset using an anisotropic Gaussian kernel. The mean map was finally generated by voxel-wise calculation of the random-effects mean of the dataset maps, weighted by the sample size, intra-dataset variability, and between-dataset heterogeneity. All analyses were conducted using the grey matter template included in AES-SDM (template sampling size of 2×2×2 mm3 voxels). The default SDM kernel size and thresholds were used (full width at half maximum [FWHM] = 20 mm, p < 0.005, peak Z value of >1, cluster size of >10). To assess heterogeneity between studies, I2 values were extracted from the meta-analytic peaks and interpreted according to (Higgins and Green, 2011). Jackknife sensitivity analyses, consisting of iteratively repeating the meta-analysis excluding one study at a time, were also conducted, for both BPD and PTSD, to examine the robustness of the main meta-analytic output. Meta-regression analyses were performed to study the effects of age and gender. Theses analyses were thresholded more conservatively (p < 0.0005) to minimize spurious findings. Publication bias was also examined by using the Egger test.

The multimodal analysis function of the AES-SDM statistical package also allows conjunction analyses to be performed, which enabled us to identify regions where both patient groups show common differences with respect to controls, while taking into account error in the estimation of the magnitude of these differences (Radua et al., 2013). The default parameters of SDM were used (peak p < 0.00025, cluster size of >10).

The main meta-analyses were performed for BPD and all the PTSD studies. Importantly, neural activity abnormalities in patients with PTSD compared to controls can appear dramatically different depending on whether individuals without trauma exposure (non-trauma exposed controls; NTC) or individuals with trauma exposure who did not develop PTSD (trauma-exposed controls; TEC) are employed as experimental controls (Disner et al., 2018). To account for such heterogeneity, the analyses were also performed using, both by merging all the PTSD studies and by performing separate analysis according to the control group. Finally, as BPD and PTSD patients represent different socio-demographic characteristics, especially for age and gender, we conducted a subgroup analysis of studies with similar sex and age distribution (more than half females and mean age less than 40 years).

## RESULTS

### Studies characteristics

#### Borderline personality disorder

Nine studies comparing patients and healthy controls (**Supplementary Table 1**) were included. These studies included a total of 235 patients and 214 healthy controls. Patients’ mean age was 28,5 years (SD=5.3) and 86.5 % were female. The mean age of healthy control participants was 30.4 years (SD=6.3), and 83.6 % were female. No information about the history of trauma and its characteristics (age and type) was available.

#### Post-traumatic stress disorder

Eighteen studies comparing patients and healthy controls (**Supplementary Table 2**) were included. These studies included a total of 433 patients and 619 healthy controls. Patients’ mean age was 39.4 years (SD=8.78) and 41.3 % were female. The mean age of healthy control participants was 40.5 years (SD=7.8), and 39.4 % were female.

### Meta-analysis

#### BPD vs healthy controls

Functional differences in BPD relative to healthy controls are shown in **Table 1** and **Figure 1**.

**Figure 1.**
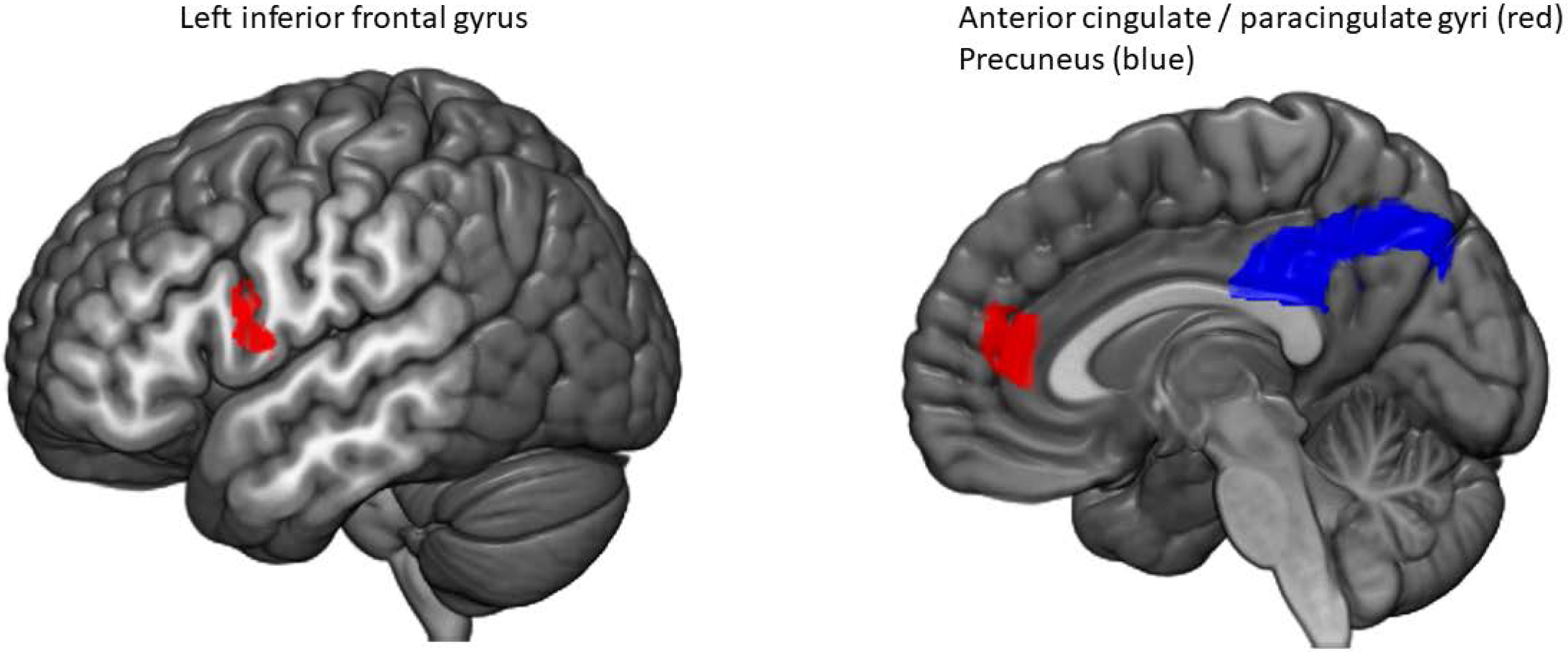
Regions of increased (red) and decreased (blue) activation at rest in patients with borderline personality disorder compared with healthy controls. Statistical maps are thresholded at p < 0.005 and k > 10.

**Table 1.** Clusters showing differences between BPD and healthy controls.

| MNI coordinates<br>(x, y, z) |  |  | SDM-Z | P | Cluster size /<br>voxels | Brodman<br>areas | Side (L/R) | Brain areas |
| --- | --- | --- | --- | --- | --- | --- | --- | --- |
| *BPD > Controls* |  |  |  |  |  |  |  |  |
| -10 | 56 | 22 | 1.647 | 0.000304401 | 805 | 32 | L+R | Anterior cingulate /<br>paracingulate gyri |
| -54 | 10 | 16 | 1.371 | 0.002392530 | 113 | 44, 48 | L | Left inferior frontal gyrus |
| *BPD < Controls* |  |  |  |  |  |  |  |  |
| 6 | -62 | 44 | -2.268 | 0.000009835 | 2043 | 7 | R | Precuneus |

We found an increased activation in patients with BPD relative to healthy controls in the anterior cingulate, paracingulate gyri and in the left inferior frontal gyrus. Reduced activation was found in the right precuneus. Heterogeneity between studies was substantial in the anterior cingulate cortex (I2 = 63 %), moderate in the left inferior gyrus (I2 = 47 %) and was not seen in the right precuneus.

#### PTSD vs healthy controls

Functional differences in PTSD relative to healthy controls are shown in **Table 2** and **Figure 2**.

**Table 2.** Clusters showing differences between PTSD and all healthy controls.

| MNI coordinates<br>(x, y, z) |  |  | SDM-Z | P | Cluster size /<br>voxels | Brodman<br>areas | Side (L/R) | Brain areas |
| --- | --- | --- | --- | --- | --- | --- | --- | --- |
| *PTSD > Controls* |  |  |  |  |  |  |  |  |
| -56 | -38 | 32 | 2.346 | 0.000039041 | 584 | 40, 48 | L | Supramarginal gyrus |
| 48 | 20 | -10 | 1.869 | 0.000604749 | 437 | 38 | R | Anterior Insula |
| -26 | -28 | -30 | 1.706 | 0.001354516 | 166 | 30 | L | Cerebellum (lobule IV / V) |
| -4 | -12 | 42 | 1.750 | 0.001094818 | 154 | 23 | L+R | Median cingulate /<br>paracingulate gyri |
| 30 | -24 | -30 | 1.689 | 0.001469672 | 80 | 20 | R | Parahippocampal gyrus |
| 38 | 42 | 38 | 1.999 | 0.000295222 | 65 | 9 | R | dIPFC |
| *PTSD < Controls* |  |  |  |  |  |  |  |  |
| 40 | -20 | 4 | -2.131 | 0.000078440 | 1625 | 48 | R | Heschl Gyrus |
| 20 | -74 | 0 | -2.202 | 0.000050783 | 164 |  | R | Lingual Gyrus |
| 52 | 22 | 38 | -1.664 | 0.001235783 | 35 | 44 | R | Inferior and Middle Frontal<br>Gyri |

**Figure 2.**
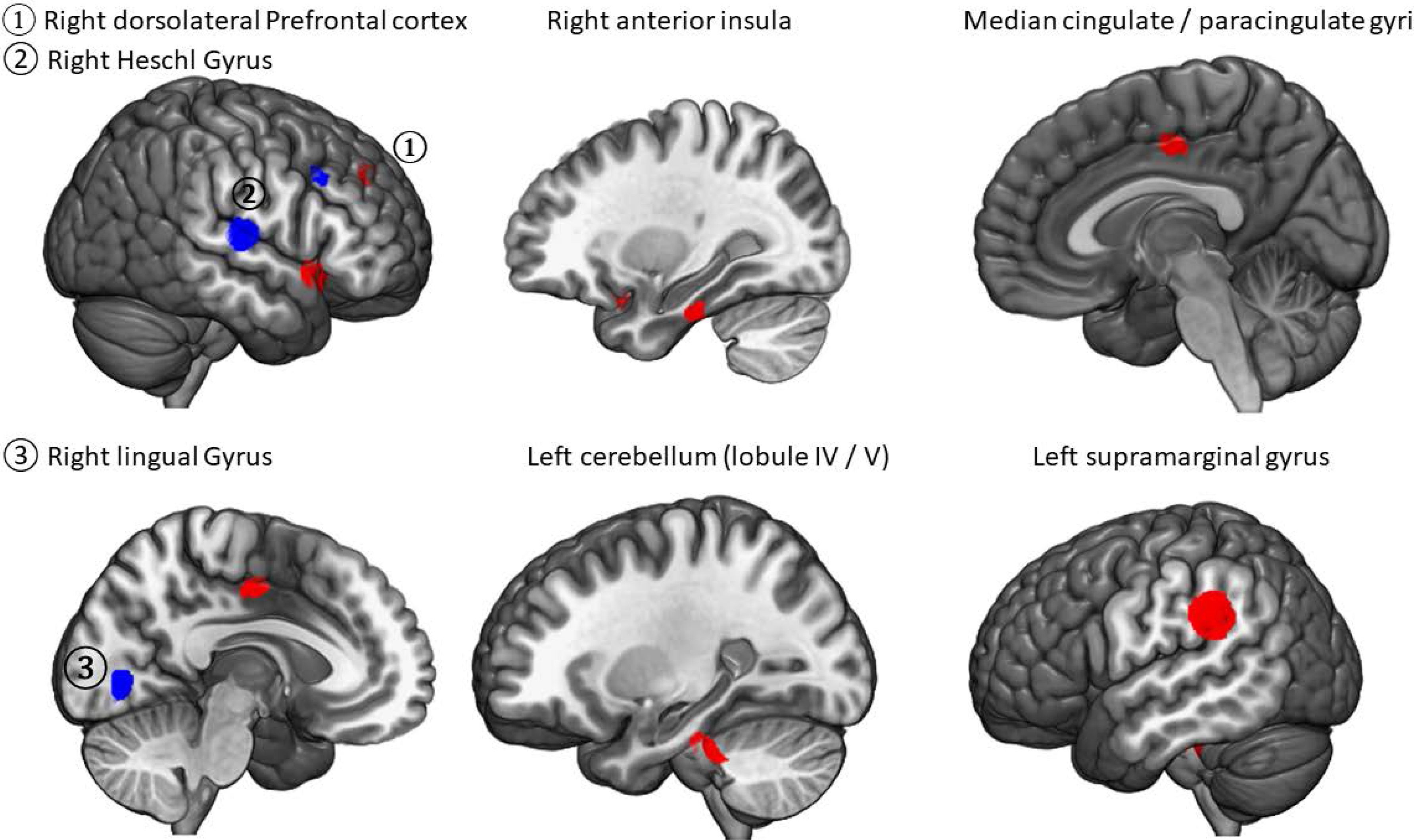
Regions of increased (red) and decreased (blue) activation at rest in patients with posttraumatic stress disorder compared with healthy controls. Statistical maps are thresholded at p < 0.005 and k > 10.

We found an increased activation in patients with PTSD relative to healthy controls in the left supramarginal, the right anterior insula, the left cerebellum, the right parahippocampal gyrus, the right dorso-lateral prefrontal cortex (dlPFC) and the median cingulate and paracingulate gyri. Reduced activations in patients with PTSD relative to healthy controls were found in the right Heschl gyrus, the right lingual gyrus and inferior and middle frontal gyri. Heterogeneity between studies was low for all the brain regions found significant (I2 < 25%). The results of the meta-analysis performed by separating the control group in TEC and NTC are summarized in **Supplementary Table 3 and 4**.

#### Reliability analysis

Jackknife sensitivity analyses showed that the main findings were highly replicable across combinations of datasets for both disorders. Indeed, the increased activity in BPD patients in anterior cingulate cortex and left inferior frontal gyrus were robust as they were significant in all but one combination. Results in the right precuneus were preserved throughout all combinations of the data sets. The activation in patients with PTSD were also highly replicable, being preserved in 16 of 24 combinations of the data sets. See **Supplementary Table 5 and 6** for details.

#### Age and gender effects

For BPD, meta-regression analysis suggested that a higher percentage of females had significantly increased activation in the left inferior frontal gyrus (MNI coordinates = −44, 42, 10; SDM = 2.697; p < 0.0001, 78 voxels) and in the anterior cingulate (MNI coordinates = 6,44,12; SDM = 2.444; p < 0.0001; 51 voxels) and a decreased activation the middle occipital gyrus (MNI coordinates = 34,−72,30; SDM = −2.739, p < 0.0001; 726 voxels). Age was associated with an increased activation in the right middle occipital gyrus (MNI coordinates = 34, −72, 32; SDM = 2.802, p < 0.0001, 758 voxels) and a decreased activation in the middle frontal gyrus (MNI coordinates = −44,44,10; SDM = −2.470; p < 0.0001; 172 voxels).

For PTSD, meta-regression analysis suggested that a higher percentage of females had significantly increased activation in the right fusiform gyrus (MNI coordinates = 20,−38,−14; SDM = 6.166; p ∼ 0; 143 voxels) and in the superior temporal gyrus (MNI coordinates = 48,12,−16; SDM = 5.629; p ∼ 0; 109 voxels) and a decrease in the left angular gyrus (MNI coordinates = −42,−62,34; SDM = −5.493; p ∼ 0; 45 voxels). Age was associated with an increased activation in the right supplementary area (MNI coordinates = 10,−2,46; SDM = 4.518; p ∼ 0; 1401 voxels) and in the right frontal superior gyrus (MNI coordinates = 22,−2,48; SDM = 2.170; p < 0.0001; 25 voxels) and a decreased activation in the left olfactory cortex (MNI coordinates = −18,4,−14; SDM = −2.018, p < 0.0001; 381voxels) and in the left lingual gyrus (MNI coordinates = −16,−98,−12; SDM = −1.546; p < 0.0001; 127 voxels).

#### Publication bias analysis

For BPD, there was no publication bias for these clusters (p-values for Egger tests > 0.15). For PTSD, only the cluster encompassing the dlPFC showed a publication bias (p = 0.003).

#### Overlap analyses

Overlap analyses (**Figure 3**) revealed a significant shared decreased activation in both BPD and PTSD groups relative to control subjects in left and right precuneus (MNI coordinates = 0, −46, 38; 606 voxels). BPD showed significant increased, but PTSD decreased activation relative to control subjects in the anterior cingulate/paracingulate gyri and in the left superior frontal gyrus (MNI coordinates = 2, 44, 20; 1205 voxels). The overlap analyses by using specifically TEC or NTC for PTSD studies are presented in **Tables 3 and 4**.

**Figure 3.**
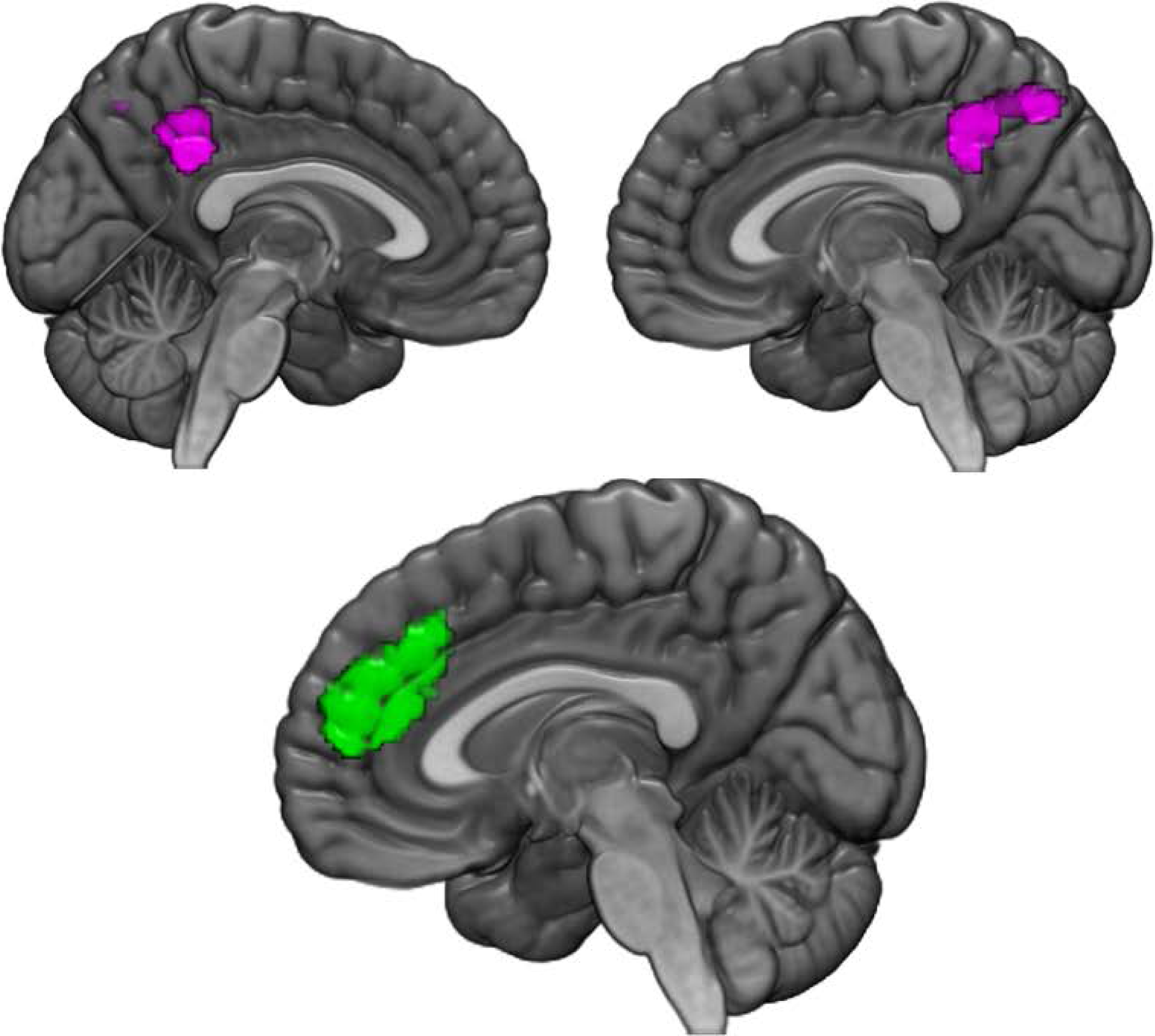
Clusters showing significant shared decreased activation in both BPD and PTSD (purple) groups relative to control subjects in left and right precuneus (MNI coordinates: 0, −46, 38; 606 voxels). Other regions (in green) were also identified where BPD showed significant increased but PTSD decreased activation relative to control subjects in the anterior cingulate/paracingulate gyri and in the left superior frontal gyrus (MNI coordinates: 2, 44, 20; 1205 voxels).

**Table 3.** Clusters showing similar activation in BPD and PTSD (trauma exposed controls, TEC).

| MNI coordinates<br>(x, y, z) |  |  | P | Cluster size /<br>voxels | Brodman<br>areas | Side (L/R) | Brain areas |
| --- | --- | --- | --- | --- | --- | --- | --- |
| *PTSD pos – bpd pos (increased activity in both ptsd and bpd)* |  |  |  |  |  |  |  |
| -46 | -6 | 4 | < 0.00025 | 649 | 48 | L | Left rolandic operculum, Left insula, Left superior temporal gyrus |
| 56 | -52 | 30 | < 0.00025 | 364 | 39,40 | R | Right angular gyrus |
| *PTSD neg – bpd neg (decreased activity in both ptsd and bpd)* |  |  |  |  |  |  |  |
| None |  |  |  |  |  |  |  |
| *PTSD pos – bpd neg (increased activity in ptsd and decreased in bpd)* |  |  |  |  |  |  |  |
| -6 | -24 | 30 | < 0.00025 | 311 | 23 | L+R | median cingulate / paracingulate gyri |
| *PTSD neg – bpd pos (decreased activity in ptsd and increased in bpd)* |  |  |  |  |  |  |  |
| -4 | 58 | 22 | < 0.00025 | 396 | 10, 32 | L | Left superior frontal gyrus, medial |
| -24 | 50 | 26 | < 0.00025 | 290 | 46 | L | Left middle frontal gyrus, Left superior frontal gyrus, dorsolateral |

**Table 4.**
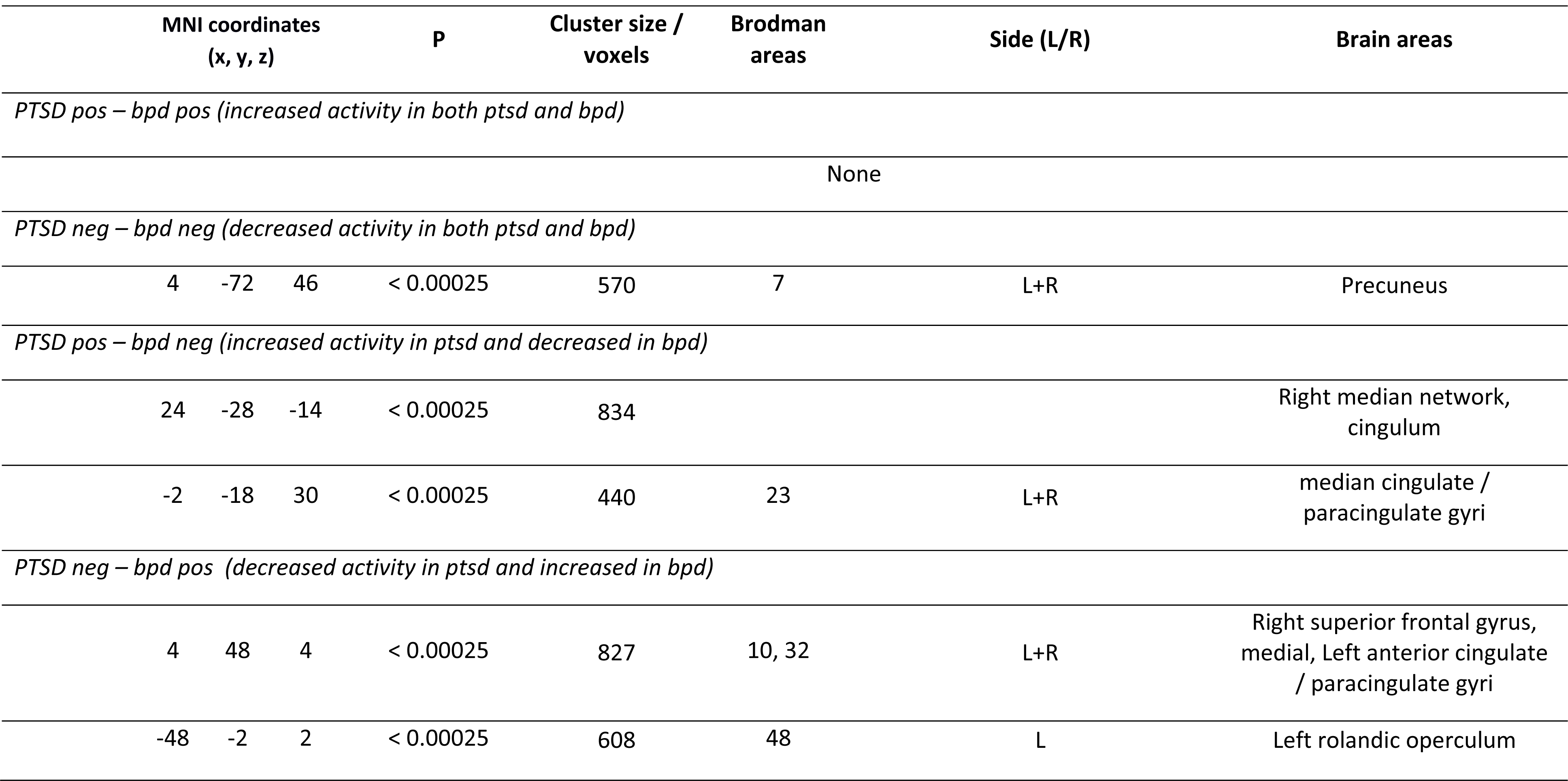
Clusters showing similar activation in BPD and PTSD (non-trauma exposed controls, NTC).

Complementary overlap analyses were also performed on a subgroup which only included studies with more than half females and a mean age of less than 40 years. Several regions were identified where BPD showed significant increased but PTSD decreased activation relative to control subjects in the anterior cingulate/paracingulate gyri and in the left superior frontal gyrus (MNI coordinates = 2, 38, 0; 506 voxels), but also in the left insula and inferior frontal gyrus (MNI coordinates = −36, 12, 6; 2169 voxels) and in the left hippocampus (MNI coordinates = −26, −8, −24; 340 voxels).

## DISCUSSION

Here, we report findings from two distinct resting-state functional meta-analyses conducted in BPD and PTSD. For BPD, we found an increased activity in patients relative to healthy controls in the anterior cingulate cortex (ACC) and in the left inferior frontal gyrus, and a reduced activity in the right precuneus, which complements previously published functional meta-analyses using the same method (Amad and Radua, 2017; Visintin et al., 2016). These results contrast with the established fronto-limbic hypothesis of BPD which postulates a decreased activity of frontal brain regions and a limbic hyperactivity. In fact, the brain regions found here may correspond to specific networks involved in pain processing (Kluetsch et al., 2012) and dissociative states (*i.e.* disintegration of perception, consciousness, identity and memory). Dissociation, characterized by subjective detachment from overwhelming emotional experience during and after a trauma, is known to be frequently found (up to two thirds) of BPD patients (Vermetten and Spiegel, 2014). Interestingly, the regions identified here have been associated in participants with PTSD while in dissociative states (Lanius et al., 2002). We also found an increased activity in the left inferior frontal gyrus a critical brain region for response inhibition and for successful implementation of inhibitory control over motor responses (Swick et al., 2008) which has also been associated with attention-deficit/hyperactivity disorder (ADHD) (van Rooij et al., 2015) another disorder characterised by impulsivity and which is often associated with BPD.

Regarding PTSD, our results are also highly consistent with a recent resting-state functional meta-analysis using the same method (Wang et al., 2016), even after adding 3 new neuroimaging studies. We found an increased activation in the anterior insula, the cerebellum, the cingulate and paracingulate gyri and the parahippocampal gyrus and a decreased activation in the Heschl and lingual gyri. When the analyses were performed with specific control groups (i.e. NTC or TEC), our results also confirmed previous studies: PTSD patients show (i) increased activation in the anterior insula and decreased activation in the dorso-medial prefrontal cortex when compared with NTC, (ii) increased activation in the ventro-medial prefrontal cortex when compared with TEC (Wang et al., 2016).

In this study, we have also directly compared BPD and PTSD to identify common and distinct patterns of functional modulations. We showed that the two conditions share similar patterns of decreased activation in left and right precuneus. We have also identified that the anterior cingulate/paracingulate gyri and left superior frontal gyrus are affected in both conditions (increased activation for BPD patients and decreased activation for PTSD subjects, relative to control subjects). These results were partially confirmed by complementary analysis on a more homogeneous subgroup regarding age and gender and when the analyses were performed using specific control groups (TEC or NTC).

Reduced activation in precuneus was found in both BPD and PTSD. The precuneus, which is part of the medial parietal cortex, plays a central role to link the limbic structures with parietal regions. This region is also involved in autobiographical memory and in self-referential processing, namely first-person perspective and experience of agency (Cavanna and Trimble, 2006). Precuneus activity has been demonstrated as related to trauma memory generalization (Hayes et al., 2011), and flashbacks (Whalley et al., 2013). Thus, alterations in the precuneus found in BPD and PTSD patients may be related to altered self-referential and memory processes, such as memory deficits, intrusions or flashbacks.

Anterior cingulate/paracingulate gyri and left superior frontal gyrus were also identified to be affected in both conditions. The anterior cingulate gyrus is part of the medial prefrontal cortex (PFC) which exerts inhibitory control over stress responses and emotional reactivity partly through connections with the amygdala. Interestingly, patients with PTSD also exhibit decreased ACC volumes (Bromis et al., 2018), that seem to be consequences of the course of PTSD rather than a pre-existing risk factor (Kasai et al., 2008). Functional imaging studies involving patients with PTSD also found decreased activation of the medial PFC in response to trauma stimuli, such as trauma scripts, combat pictures and sounds, trauma-unrelated negative narratives, fearful faces, emotional stroop (for review see (Sherin and Nemeroff, 2011)). Even if there are discordant findings, reduced activation of the medial PFC was associated with PTSD symptom severity in several studies. Successful SSRI treatment has also been shown to restore medial PFC activation patterns (Shin et al., 2006). Regarding BPD, higher levels of glutamate in the ACC were found in patients when compared with healthy controls, and there are positive correlations between glutamate levels and impulsivity (Hoerst et al., 2010). It has also been shown that the hyperactivation of ACC was correlated with hippocampal activity, abnormal functioning of the memory system, poorer performance in retrieval tasks related to traumatic or emotionally significant memories (Mensebach et al., 2009; Sala et al., 2009; Schmahl et al., 2004).

We believe that our results are in line with the hypothesis that the conceptual interface between BPD and PTSD could be highlighted by the ‘*age-dependent neuroplasticity*’ framework (Amad et al., 2016) which postulates that the same experience can differentially affect the brain plasticity depending on one’s age (Kolb and Gibb, 2014). Indeed, neuroplasticity (*i.e.* the ability of the nervous system to respond to intrinsic or extrinsic stimuli) is a crucial process involved in normal development and maturation of the brain but also in response and adaptation of the brain to the external environment or to pathological processes (Cramer et al., 2011). For example, it has been shown that the neuroplastic changes following a cerebral injury can vary greatly depending on the age of the patients (Kolb and Gibb, 2014). With respect to the effects of trauma on the brain, there is some evidence for the implication of an *age-dependent neuroplasticity* process, where the effects of stressors occurring early in development appear to be stronger and more persistent in comparison with stressors occurring during adulthood (Hoffmann and Spengler, 2014). In this perspective, it has then been proposed that trauma during childhood could favour the development of BPD (Amad et al., 2014a) while trauma in adulthood could increase the risk of PTSD (Amad et al., 2016). Moreover, it has been shown that the stress axis regulation appears to be different according to the age of trauma onset (Kuhlman et al., 2015) and it is also well known that the stress axis has different windows of regulation after a trauma (Shalev et al., 2008). However, this hypothesis could not be tested since no BPD studies included data about the trauma history of these patients.

One of the main limitations of this study corresponds to the very different socio-demographic characteristics of BPD and PTSD patients included, especially for age and gender as highlighted by the meta-regression analysis. Accordingly, we conducted a complementary analysis focusing on studies with more than half females and a mean age of less than 40 years. This complementary analysis partially confirmed our main results as we found the same pattern of functional activation in the ACC and in the left superior frontal gyrus between BPD and PTSD. Another possible limitation is the use of different statistical approaches such as independent component analysis, regional homogeneity, etc. to assess resting-state in the different studies included (Kühn and Gallinat, 2013). As soon as more resting-state studies on BPD and PTSD patients are published, new analysis should be performed including studies with the same statistical approach and future research will also need to confirm our results. Cautious interpretation is also needed because BPD has a potentially high clinical heterogeneity and has been associated with many comorbidities, including mood disorders, anxiety disorders, and PTSD. Unfortunately, data presented in the different studies included in these meta-analyses did not allow to include these potential confounding factors in our analyses. The results therefore can hardly be attributed solely to BPD or PTSD. In the same way, no sufficient information about medication was available to perform complementary analysis. Future studies should focus on the clinical assessment of carefully selected BPD and PTSD patients to explore specific dimensions and refined phenotypes (e.g., type of trauma, psychotic features or impulsivity) to improve the comprehension of the similarities between these disorders.

One of the most frequent criticism when considering BPD as a trauma spectrum disorder is that the existence of trauma is not necessary or sufficient to explain the development of the disorder (Lewis and Grenyer, 2009). However, it is now well established that gene-environment interactions play a role in the development of both BPD (Amad et al., 2014a) and PTSD (Koenen et al., 2008). Moreover, clinical profiles of BPD patients appear to differ according to the severity of childhood trauma (Zanarini et al., 2002).

In conclusion, this study sheds new light on the neurobiology of BPD, which has been little investigated despite its high prevalence (0.5–5.9% of the general population) and severity (Amad et al., 2014b) whilst also highlighting the functional similarities between BPD and PTSD. Our findings help reinforce the hypothesis that BPD shares conceptual and phenomenological similarities with PTSD. One difference between these disorders could be the age at which trauma occurs, differentially affecting the resting brain function and thus the psychiatric symptoms (Amad et al., 2016).

## Conflict of interests

The authors declare no potential conflict of interest.

## Contributors

AA, JR, TF participated in the conception and design of the study; AA and TF participated in the acquisition of data; AA, JR, TF performed the analyses; AA and TF wrote the first draft of the manuscript. All authors participated in the writing and revision of the successive drafts of the manuscript and approved the final version.

## Role of the Funding source

No funding source to declare.

## Acknowledgements

No Acknowledgement

