## Supplementary figures and tables for "Similarities between Borderline Personality Disorder and Post traumatic Stress Disorder: evidence from Resting-State Meta-Analysis"

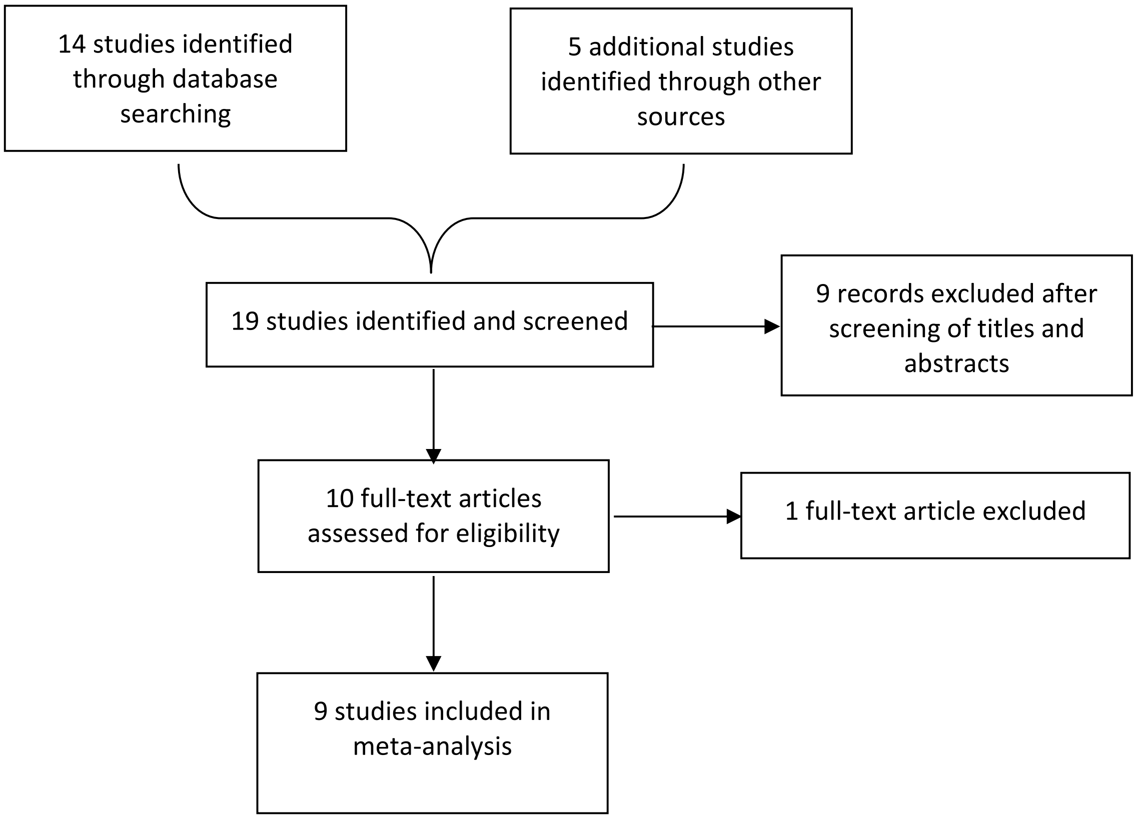


**Supplementary Figure 1**. Flowchart of selection of BPD studies.

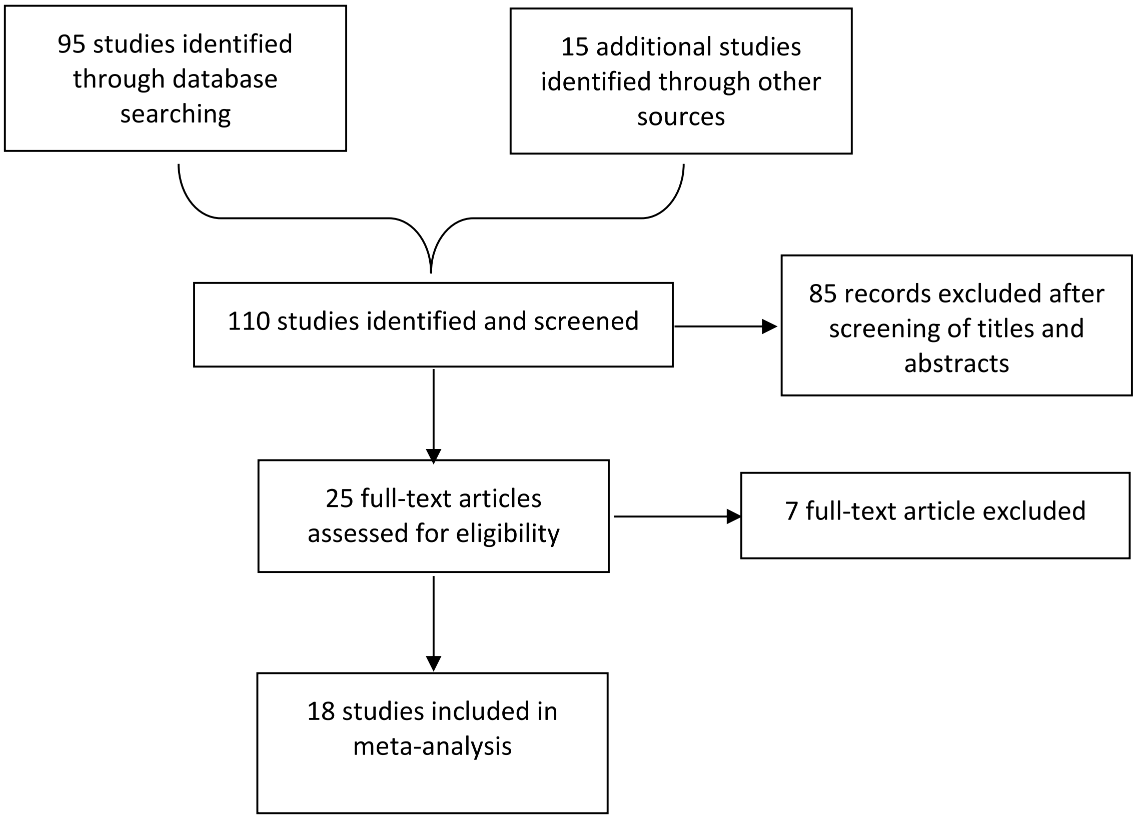


**Supplementary Figure 2**. Flowchart of selection of PTSD studies.

| Study | Imaging modality and method of analysis | Patients | | Controls | | Diagnosis criteria | Reported contrast | Medication status | Main current comorbidities | Statistical Threshold | Main findings in BPD patients in comparison with controls |
|---|---|---|---|---|---|---|---|---|---|---|---|
| | | *n (F)* | *Age (mean +/- SD)* | *n (F)* | *Age (mean +/- SD)* | | | | | | |
| Juengling et al. (Juengling et al., 2003) | FDG PET | 12(12) | 25(±4) | 12(12) | 30(±9) | DSM IV | • ctrl>bpd<br>• ctrl<bpd | Unmedicated for 4 weeks | • Anxiety disorders (83%)<br>• Bulimia nervosa (17%)<br>• Dysthymia (33%)<br>• PTSD (25%) | Corrected | • Frontal and prefrontal hypermetabolism<br>• Hypometabolism in hippocampus and cuneus |
| Soloff et al. (Soloff et al., 2003) | FDG PET | 13(13) | 25.2(±7.1) | 9(9) | 27.4(±6.4) | DSM III-R | • ctrl>bpd<br>• ctrl<bpd | Unmedicated for at least 3 months | • Alcohol abuse (31%)<br>• Dysthymia (54%)<br>• Substance abuse (15%) | Corrected | • Bilateral medial orbital frontal hypometabolism |
| Lange et al. (Lange et al., 2005) | FDG PET | 17(17) | 32(±4) | 9(9) | 33(±6) | DSM IV | • ctrl>bpd<br>• ctrl<bpd | 5 unmedicated and 12 medicated patients | • Anxiety disorders (89%)<br>• Bulimia nervosa (18%)<br>• MDD (71%)<br>• PTSD (35%) | Corrected | • Hypometabolism in the right temporal pole/anterior fusiform gyrus and in the left precuneus and posterior cingulate cortex |
| Lei et al. (Lei et al., 2017) | fMRI/ALFF & ReHo | 40(20) | 25.2(±3.2) | 35(15) | 24.8(±1.2) | DSM-IV | • ctrl>bpd<br>• ctrl<bpd | Unmedicated | • Unknown | Corrected | • Decreased ALFF and ReHo both in the right posterior cingulate cortex (PCC) and adjacent precuneus |
| Salavert et al. (Salavert et al., 2011) | FDG PET | 8(6) | 35.5(±9.27) | 8(5) | 32(±7.86) | DSM-III-R | • ctrl>bpd<br>• ctrl<bpd | Medicated | • Unknown | Corrected | • Hypermetabolism in motor cortex, medial and anterior cingulus, occipital lobe, temporal pole, left superior parietal gyrus and right superior frontal gyrus<br>• Hypometabolism in frontal lobe |
| Wolf et al. (Wolf et al., 2011) | fMRI/ICA | 17(17) | 27.2(±8) | 17(17) | 28.6(±7.3) | DSM IV | • ctrl>bpd<br>• ctrl<bpd | Medicated | • Anxiety disorders (6%)<br>• Eating disorders (24%)<br>• MDD (53%) | Corrected | • Increased FC in the left prefrontal cortex and the left insula<br>• Decreased FC in the left cuneus, the left inferior parietal lobule and the right middle temporal gyrus |
| Doll et al. (Doll et al., 2013) | fMRI/ICA | 14(13) | 30.4 | 16(15) | 34.0 | DSM IV | • ctrl>bpd<br>• ctrl<bpd | Medicated | • PTSD (14%)<br>• Alcohol abuse (36%)<br>• MDD (14%)<br>• Substance abuse (21%) | Corrected | • Increased FC in anterior and posterior cingulate cortex, medial frontal gyrus and parietal lobe<br>• Decreased FC in right hippocampus and left superior frontal gyrus. |
| Das et al. 2014 (Das et al., 2014) | fMRI/ICA | 14 (14) | 32 (±7.86) | 13 (13) | 31.75(±11.07) | DSM IV | None | Medicated | • Unknown | Uncorrected | • Functional connectivity strength between social salience network and right fronto-parietal network was significantly reduced in BPD compared to controls |

| | | | | | | | | | | | |
|---|---|---|---|---|---|---|---|---|---|---|---|
| Salvador et al. (Salvador et al., 2014) | fMRI/ALFF | 60(60) | 32.12(±7.16) | 60(60) | 33.73(±12.8) | DSM IV | • ctrl>bpd<br>• ctrl<bpd | Medicated | • Unknown | Corrected | • Increased amplitudes in the left hippocampus and amygdala and in the left putamen<br>• Reduces amplitudes in the occipital lobes, in the right precuneus, and in the dorsalposterior cingulate cortex. |

**Supplementary table 1:** Studies included in the meta-analysis. ALFF, amplitude of low frequency fluctuations; BPD, borderline personality disorder; CTRL: controls; DSM: Diagnostic and Statistical Manual of Mental Disorders; FC functional connectivity FDG PET, fludeoxyglucose Positron emission tomography; fMRI, functional MRI; ICA, independent component analysis; MDD, major depressive disorder, PTSD, post-traumatic stress disorder.

| Study | Imaging modality and method of analysis | Trauma type | Patients PTSD n (F) | | | Trauma exposed controls n (F) | | Non trauma exposed controls n (F) | | Diagnosis criteria | Reported contrast | Medication status | Main current comorbidities | Statistical Threshold | Main findings in PTSD patients in comparison with controls |
|---|---|---|---|---|---|---|---|---|---|---|---|---|---|---|---|
| | | | *n (F)* | *Age (mean±SD)* | *CAPS (mean±SD)* | *n (F)* | *Age (mean± SD)* | *n (F)* | *Age (mean±S D)* | | | | | | |
| Bing et al. (Bing et al., 2013) | fMRI/ALFF | Motor vehicle accident | 20(7) | 32.92 (±8.48)/ | 52.33(±9.44) | - | - | 20(6) | 31.53 (±7.43) | DSM-IV | • NTC vs PTSD | Unmedicated for at least 2 months | Depressive disorder (n=3) | corrected | • Increased ALFF in the left medial prefrontal cortex, the anterior cingulated and the right cerebellum. |
| Bonne et al. (Bonne et al., 2003) | SPECT/rCBF | Civilian traumatic events (most motor vehicle accidents) | 11(7) | 34(±9) | 57.9(±17.85) | 17(9° | 35(±8) | 11(6) | 33(±12) | DSM-IV | • NTC vs PTSD<br>• TEC vs PTSD | Unmedicated | None | uncorrected | • rCBF in the cerebellum was higher in PTSD disorder than in both control groups. rCBF in right precentral, superior temporal, and fusiform gyri in PTSD was higher than in healthy controls. |
| Chung et al. (Chung et al., 2006) | SPECT/rCBF | Civilian traumatic events (motor vehicle accidents n=18) | 23(10) | 43 | 87.9(±12.7) | - | | 64(30) | 43 | DSM-IV | • NTC vs PTSD | Unmedicated for 2 weeks | Unknown | corrected | • PTSD patients exhibited increased cerebral blood perfusion in limbic regions and decreased perfusion in the superior frontal gyrus and parietal and temporal regions |
| Ke et al. (Ke et al., 2017) | fMRI/ReHo | Typhoon | 27(20) | 48.4(±10.3) | 78.2(±19.3) | 33(26) | 48.5(±7.5) | 30(23) | 49.9(±6.1) | DSM-IV | • NTC vs PTSD<br>• TEC vs PTSD | Unmedicated | None | corrected | • The PTSD group showed ReHo changes in multiple regions, including the amygdala, parahippocampal gyrus, and prefrontal cortex relative to both control groups. Compared with healthy controls, typhoon survivors had increased ReHo in the insula/inferior frontal gyrus, middle and dorsal anterior cingulate cortex (MCC/dACC), as well as enhanced negative FC between the MCC/dACC and posterior cingulate cortex (PCC)/precuneus. The typhoon-exposed control group exhibited higher ReHo in the PCC/precuneus than the |

| | | | | | | | | | | | | | | | |
|---|---|---|---|---|---|---|---|---|---|---|---|---|---|---|---|
| | | | | | | | | | | | | | | | PTSD and healthy control groups. |
| Kim et al. (Kim et al., 2007) | SPECT/rCBF | Subway fire | 19(13) | 26.6(±7.2) | 71 | - | - | 19(7) | 31.6(±5.5) | DSM-IV | • NTC vs PTSD | Unknown | Depressive disorders and anxiety disorders | corrected | • PTSD patients showed a decreased rCBF in the right thalamus and an increased rCBF in the right superior parietal lobe relative to comparison subjects |
| Kim et al. (Kim et al., 2012) | SPECT/rCBF PET/rCMRglu | Sexual assaults | 12(12) | 35.9(±13.8) | NR | - | - | 10(10) (SPECT) 15(15) (PET) | 37.2(±10.4) 38.4(±12.1) | DSM-IV | • NTC vs PTSD (SPECT) • NTC vs PTSD (PET) | Venlafaxine (75–225 mg/day) for a mean period of 5.2 months | Unknown | uncorrected | • The PTSD patients showed significant relative decreases in perfusion in the left hippocampus and in the basal ganglia compared with the control group. The PTSD group also had significantly lower cerebral glucosemetabolic activity in the left hippocampus and the superior temporal and precentral gyri than in the control group. |
| Li et al. (Li et al., 2013) | ASL-fMRI/rCBF | Coal mining flood disaster | 10(0) | NR | NR | 10(0) | NR | - | - | DSM-IV | • TEC vs PTSD | Unknown | Unknown | uncorrected | • Decreased regional CBF in the right middle temporal gyrus, lingual gyrus, and postcentral gyrus was detected in the PTSD patients. |
| Shang et al. (Shang et al., 2014) | fMRI/ICA | Earthquake | 18(14) | 43.33(±8.04) | 63.94(±13.53) | 20(9) | 40.3(±9.3) | - | - | DSM-IV | • TEC vs PTSD | Unmedicated | Unknown | corrected | • PTSD patients displayed both increased and decreased functional connectivity within the Salience network (SN), central executive network (CEN), default mode network (DMN), somato-motor network (SMN), auditory network (AN), and visual network (VN). |
| Schuff et al. (Schuff et al., 2011) | ASL-fMRI/rCBF | Combat | 17(0) | 45(±14) | 62(±13) | 15(0) | 37(±13) | - | - | DSM-IV | • TEC vs PTSD | Unknown | Unknown | uncorrected | • Subjects with PTSD had increased rCBF in primarily right parietal |

| | | | | | | | | | | | | | | | |
|---|---|---|---|---|---|---|---|---|---|---|---|---|---|---|---|
| | | | | | | | | | | | | | | | and superior temporal cortices. |
| Shin et al. (Shin et al., 2009) | PET/rCMRglu | Combat | 14(0) | 57.8(±2.8) | 66(±25.8) | 19(0) | 57.1(±2.2) | 14 (co-twins NTC) 19 NTC | 57.8(±2.8) 57.1(±2.2) | DSM-IV | • TEC vs PTSD • NTC vs PTSD | Antidepressants (n=13) Benzodiazepines (n=2) | Depressive disorders, anxiety disorders, substance use disorders | uncorrected | • Veterans with PTSD and their co-twins had significantly higher resting rCMRglu in dorsal anterior cingulate/mid cingulate cortex (dACC/MCC) compared to non-PTSD veterans and their co-twins. |
| Yan et al. (Yan et al., 2013) | fMRI/ALFF | Combat | 52(0) | 33.18 (±7.60) | 66.75(±2.69) | 52(0) | 33.57 (±8.98) | - | - | DSM-IV | • TEC vs PTSD | Unknown | Unknown | corrected | • PTSD subjects showed increased spontaneous activity in the amygdala, ventral anterior cingulate cortex, insula, and orbital frontal cortex, as well as decreased spontaneous activity in the precuneus, dorsal lateral prefrontal cortex and thalamus |
| Yin et al. (Yin et al., 2011a) | fMRI/ReHo | Earthquake | 54(NR) | NR | 64.09(±9.68) | 72(NR) | NR | - | - | DSM-IV | • TEC vs PTSD | Unknown | • None | corrected | • PTSD patients presented enhanced ReHo in the left inferior parietal lobule and right superior frontal gyrus, and reduced ReHo in the right middle temporal gyrus and lingual gyrus, relative to traumatized individuals without PTSD. |
| Yin et al. (Yin et al., 2011b) | fMRI/ALFF | Earthquake | 54(39) | 42 | 64.09(±9.68. | 72(50) | 42 | - | | DSM-IV | • TEC vs PTSD | Unmedicated | • Unknown | corrected | • PTSD patients showed decreased ALFF values in right lingual gyrus, cuneus, middle occipital gyrus, insula, and cerebellum, and increased ALFF values in right medial and middle frontal gyri, relative to traumatized individuals without PTSD |
| Zhang et al. (Zhang et al., 2015) | fMRI/ICA | Motor vehicle accidents | 20(7) | 32.92(±8.48) | 52.33(±9.44) | - | - | 20(6) | 31.53(±7.43) | DSM-IV | • NTC vs PTSD | Unmedicated for 2 months | • Unknown | corrected | • Compared with HCs, the PTSD patients exhibited significantly decreased Network connectivity within the anterior default mode network, |

| | | | | | | | | | | | | | | | |
|---|---|---|---|---|---|---|---|---|---|---|---|---|---|---|---|
| | | | | | | | | | | | | | | | posterior default mode network (pDMN), saliencenetwork(SN),sensory-motornetwork,andauditorynetwork. |
| Zhe et al. (Zhe et al., 2016) | fMRI/rCBF | Coal mine flood | 30(13) | 33.3(±9.8) | 49.9(±27.4) | - | - | 36(16) | 34.4(±7.8) | DSM-IV | • NTC vs PTSD | Unmedicated | • None | corrected | • PTSD symptom severity was associated with diminished cerebral blood flow in the right insular cortex and right orbital medial frontal gyrus. |
| Zhu et al. (Zhu et al., 2014) | fMRI/ALFF | Earthquake | 17(12) | 44.41(±8.44) | 60.88(±14.29) | 20(11) | 40.35(±9.43) | - | - | DSM-IV | • TEC vs PTSD | Unmedicated | • None | corrected | • Compared to traumatized controls, the PTSDgroup showed significantly altered ALFF in many emotion-related brain regions, such as the medial anterior cingulate cortex (MACC), dorsolateral prefrontal cortex (DLPFC), insular (IC), middle temporal gyrus (MTG), and ventral posterior cingulate cortex (VPCC). |
| Zhu et al. (Zhu et al., 2015) | fMRI/ALFF | Earthquake | 21(17) | 46.76(±5.81) | 68.76(±16.34) | 17(12) | 43.24(±10.62) | - | - | DSM-IV | • TEC vs PTSD | Unmedicated | • Depressive disorders, anxiety disorders | corrected | • Hyperactive function of visual cortex was observed in PTSD patients. |
| Zhong et al. (Zhong et al., 2015) | fMRI/ReHo | Civilian traumatic events (motor vehicle accidents n=7) | 14(8) | 31.3(±9.24) | 67.95(±10.09) | - | - | 14(8) | 28.5(±6.3) | DSM-IV | • NTC vs PTSD | Unmedicated | • Depressive disorder (n=7) | corrected | • Compared with the normal controls, PTSD patients showed increased local coherence in subcortical regions, including amygdala, hippocampus, thalamus, and putamen, and decreased local coherence in cortical regions, including medial prefrontal cortex and dorsolateral prefrontal cortex. |

**Supplementary table 2:** Studies included in the meta-analysis. ALFF, amplitude of low frequency fluctuations; CTRL: controls; DSM: Diagnostic and Statistical Manual of Mental Disorders; FC functional connectivity FDG PET, fludeoxyglucose Positron emission tomography; fMRI, functional MRI; ICA, independent component analysis; MDD, major depressive disorder, NTC, non-trauma exposed control; TEC, trauma exposed control; PTSD, post-traumatic stress disorder.

| | MNI coordinates (x, y, z) | | | SDM-Z | P | Cluster size / voxels | Brodman areas | Side (L/R) | Brain areas |
|---|---|---|---|---|---|---|---|---|---|
| *PTSD > TEC* | | | | | | | | | |
| | -52 | -32 | 32 | 2.561 | 0.000011563 | 1020 | | L | Supramarginal Gyrus |
| | 0 | 50 | -14 | 2.180 | 0.000144601 | 731 | 11 | L | vmPFC |
| | 30 | 48 | 32 | 2.609 | 0.000008643 | 386 | 46 | R | dlPFC |
| | 58 | -54 | 24 | 1.665 | 0.001578987 | 296 | 22 | R | Angular gyrus |
| *PTSD < TEC* | | | | | | | | | |
| | 46 | -18 | 2 | -1.759 | 0.000753582 | 1111 | | R | Heschl Gyrus |
| | 20 | -74 | 0 | -2.897 | 0.000001013 | 322 | | R | Lingual Gyrus |
| | 28 | -68 | -18 | -1.629 | 0.001604378 | 80 | 19 | R | Cerebellum (lobule VI) |
| | 58 | 2 | -26 | -1.656 | 0.001384020 | 74 | 21 | R | Temporo-parietal Junction |
| | 50 | 22 | 42 | -1.676 | 0.001234770 | 46 | 44 | R | Middle Frontal Gyrus |
| | -54 | 18 | 30 | -1.536 | 0.002774477 | 17 | 44 | L | Inferior Frontal Gyrus |

**Supplementary Table 3.** Clusters showing differences between PTSD and trauma-exposed controls (TEC).

| MNI coordinates (x, y, z) | | | SDM-Z | P | Cluster size / voxels | Brodman areas | Side (L/R) | Brain areas |
|---|---|---|---|---|---|---|---|---|
| *PTSD > NTC* | | | | | | | | |
| 28 | -32 | -24 | 2.246 | 0.000024676 | 1128 | BA 20 | Right | Parahippocampal Gyrus |
| 46 | 20 | -6 | 2.111 | 0.000076294 | 703 | | Right | Anterior Insula |
| -4 | -14 | 44 | 1.627 | 0.001889586 | 144 | BA 23 | Left | Median Cingulate / Paracingulate Gyri |
| -20 | -26 | -30 | 1.587 | 0.002381325 | 27 | 30 | Left | Cerebellum (lobule IV / V) |
| 0 | 14 | -24 | 1.518 | 0.003511727 | 12 | 11 | Left | Gyrus Rectus |
| *PTSD < NTC* | | | | | | | | |
| 4 | 54 | -6 | -1.664 | 0.001134872 | 236 | 10 | Right | vmPFC |
| 46 | -76 | 28 | -1.732 | 0.000762343 | 144 | 39 | R | Precuneus |

**Supplementary Table 4:** Clusters showing differences between PTSD and non-trauma exposed controls.

| | Hyperactivation | | Hypoactivation |
|---|---|---|---|
| **Discarded studies** | **Anterior cingulate / paracingulate gyri** | **Left inferior frontal gyrus** | **Precuneus** |
| **Doll** | Yes | Yes | Yes |
| **Juengling** | No | Yes | Yes |
| **Lei** | Yes | Yes | Yes |
| **Lange** | Yes | Yes | Yes |
| **Salavert** | Yes | Yes | Yes |
| **Salvador** | Yes | Yes | Yes |
| **Soloff** | Yes | Yes | Yes |
| **Wolf** | Yes | Yes | Yes |

**Supplementary Table 5:** Jackknife analyses of studies in the BPD vs. controls meta-analysis.

| | Hyperactivation | | | | | | Hypoactivation | | |
|---|---|---|---|---|---|---|---|---|---|
| **Discarded studies** | **Left Supramarginal gyrus** | **Right Anterior Insula** | **Left Cerebellum (lobule IV / V)** | **Median cingulate / paracingulate gyri** | **Right Parahippocampal gyrus** | **Right dlPFC** | **Right Heschl Gyrus** | **Right Lingual Gyrus** | **Right Inferior and Middle Frontal Gyri** |
| **Bing** | Yes | Yes | Yes | Yes | Yes | Yes | Yes | Yes | Yes |
| **Bonne (NTC)** | Yes | Yes | No | Yes | Yes | Yes | Yes | Yes | Yes |
| **Bonne (TEC)** | Yes | Yes | No | Yes | Yes | Yes | Yes | Yes | Yes |
| **Chung** | Yes | No | Yes | Yes | No | Yes | Yes | Yes | Yes |
| **Ke (NTC)** | Yes | No | Yes | No | No | Yes | Yes | Yes | Yes |
| **Ke (TEC)** | Yes | Yes | Yes | Yes | Yes | Yes | Yes | Yes | Yes |
| **Kim 2006** | Yes | Yes | Yes | Yes | Yes | Yes | Yes | Yes | Yes |
| **Kim2012 (PET)** | Yes | Yes | Yes | Yes | Yes | Yes | Yes | Yes | Yes |
| **Kim2012 (SPECT)** | Yes | Yes | Yes | Yes | Yes | Yes | Yes | Yes | Yes |
| **Li** | Yes | Yes | Yes | Yes | Yes | Yes | Yes | Yes | Yes |
| **Li (rcbf)** | Yes | Yes | Yes | Yes | Yes | Yes | Yes | Yes | Yes |
| **Shang** | Yes | Yes | Yes | Yes | Yes | Yes | Yes | Yes | Yes |
| **Schuff** | Yes | Yes | Yes | Yes | Yes | Yes | Yes | Yes | Yes |
| **Shin (TEC)** | Yes | Yes | Yes | Yes | Yes | Yes | Yes | Yes | Yes |
| **Shin (NTC)** | Yes | Yes | Yes | Yes | Yes | Yes | Yes | Yes | Yes |
| **Shin (twin NTC)** | Yes | Yes | Yes | Yes | Yes | Yes | Yes | Yes | Yes |
| **Yan** | Yes | Yes | Yes | Yes | Yes | Yes | Yes | Yes | No |
| **Yin 2011** | Yes | Yes | Yes | Yes | No | No | No | Yes | Yes |
| **Yin 2012** | Yes | Yes | Yes | Yes | Yes | No | Yes | No | Yes |
| **Zhang** | Yes | Yes | Yes | Yes | Yes | Yes | Yes | Yes | Yes |
| **Zhe** | Yes | Yes | Yes | Yes | Yes | Yes | No | Yes | Yes |
| **Zhu 2014** | Yes | Yes | Yes | Yes | Yes | Yes | Yes | Yes | Yes |
| **Zhu2015** | Yes | Yes | Yes | Yes | Yes | Yes | Yes | Yes | Yes |
| **Zhong** | Yes | Yes | Yes | Yes | Yes | Yes | Yes | Yes | Yes |

**Supplementary Table 6:** Jackknife analyses of studies in the PTSD vs. controls meta-analysis.
